# CAFE: An Integrated Web App for High-Dimensional Analysis and Visualization in Spectral Flow Cytometry

**DOI:** 10.1101/2024.12.03.626714

**Authors:** Md Hasanul Banna Siam, Md Akkas Ali, Donald Vardaman, Satwik Acharyya, Mallikarjun Patil, Daniel J. Tyrrell

**Affiliations:** Department of Pathology, Heersink School of Medicine, University of Alabama at Birmingham, Birmingham, AL, 35205 USA; Department of Biostatistics, University of Alabama at Birmingham, Birmingham, AL, 35233 USA; Department of Biomedical Informatics and Data Science, University of Alabama at Birmingham, Birmingham, AL, 35294 USA

**Keywords:** Cytometry, Bioinformatics, Protocol, Leiden

## Abstract

Spectral flow cytometry provides greater insights into cellular heterogeneity by simultaneous measurement of up to 50 markers. However, analyzing such high-dimensional (HD) data is complex through traditional manual gating strategy. To address this gap, we developed CAFE as an open-source Python-based web application with a graphical user interface. Built with Streamlit, CAFE incorporates libraries such as Scanpy for single-cell analysis, Pandas and PyArrow for efficient data handling, and Matplotlib, Seaborn, Plotly for creating customizable figures. Its robust toolset includes density-based down-sampling, dimensionality reduction, batch correction, Leiden-based clustering, cluster merging and annotation. Using CAFE, we demonstrated analysis of a human PBMC dataset of 350,000 cells identifying 16 distinct cell clusters. CAFE can generate publication-ready figures in real time via interactive slider controls and dropdown menus, eliminating the need for coding expertise and making HD data analysis accessible to all. CAFE is licensed under MIT and is freely available at https://github.com/mhbsiam/cafe.

## Introduction

Flow cytometry is a widely used technique in immunology to identify and quantify immune cells based on specific surface markers^1^. The development of spectral flow cytometry (SFCM) has further expanded immunophenotyping capabilities allowing the simultaneous analysis of a greater number of parameters through the complete emission spectra of fluorophores^1^. Compared to conventional flow cytometry, SFCM uses spectral unmixing algorithms to deconvolute the overlapping signals and achieves enhanced resolution and sensitivity to distinguish between different cell populations^2^. SFCM can incorporate broader range of antibodies with up to 50 colors in a single panel improving upon the conventional FCM where the number of parameters is limited by the instrument constraints^3^. Incorporating more parameters substantially increases the complexity in gating strategy which largely relies on established convention and prior knowledge^4,5^. Additional gating steps and combinations of markers used to subset cells complicate the interpretation of such high-dimensional data. Several clustering methods are available to identify cell populations such as FlowSOM^6^, xShift^7^, SPADE^8^ and Phenograph^9^. SPADE and FlowSOM utilize hierarchical clustering with the latter employing self-organizing maps (SOMs) to cluster cells, whereas xShift detects clusters based on shifts in local cell density^6–8^. Phenograph, by contrast, constructs a K-nearest neighbor graph and applies the Louvain algorithm to identify cell clusters, but Louvain can produce poorly connected or disconnected communities^9,10^.

Recently, the Leiden clustering algorithm has emerged as a faster and more accurate alternative to improve community detection in networks^10^. Single-cell RNA sequencing (scRNA-seq) tools: Seurat^11^ (R) and Scanpy^12^ (Python) have integrated Leiden algorithms for community detection. However, running Leiden within Seurat resulted in drawbacks including higher memory usage, longer calculation time and random crashes in docker containers^13^. Scanpy resolves these issues, and unlike Seurat, Scanpy improves visualization quality by using consistent KNN and SNN graphs for both clustering and uniform manifold approximation and projection (UMAP) ^13,14^. In February 2020, Phenograph version 1.5.3 was released, which incorporated an option to use Leiden for clustering; however, the default parameter is set to Louvain through the latest release (v.1.5.7). In our previous work, we showed that the use of Leiden algorithm in community detection for SFCM data provides superior result to Phenograph (Louvain), FlowSOM, and xShift^15^. Currently, there is a scarcity of open-source tools to utilize Leiden algorithm for SCFM data analysis^16^.

Here we present CAFE, Cell Analyzer for Flow Experiment, a user-friendly web application developed in Python that works across Windows, MacOS, and Linux. The app is lightweight and can perform high-dimensional SFCM data analysis using a standard computing machine (i.e., Apple M1 chip with 16gb RAM), and it provides the flexibility to be deployed on HPC clusters for enhanced scalability. Once installed, the tool runs entirely offline and does not require an active internet connection to load files. This also enables users to maintain compliance with data security, especially with protected health information (PHI) when analyzing patient samples. CAFE can be used to process data, reduce dimension, batch correction, run Leiden clustering, perform statistics and generate a wide range of figures. Figures can be adjusted and viewed within the tool in real time. Additionally, the tool offers Kernel Density Estimation (KDE)-based data downsampling, advanced clustering with predefined markers, cluster quality evaluation, merging subclusters into metaclusters, and cell type annotation. Designed as an open-source interactive data analysis platform, CAFE enables biologists with no-coding experience to analyze SFCM data and create publication quality visualization with customizable parameters. CAFE is freely available to download at: https://github.com/mhbsiam/cafe

## Methods

### Implementation

The CAFE webtool was developed using Python programming language due to its compatibility with Scanpy library^12^. Figure 1 illustrates the components and workflows of CAFE. Streamlit (streamlit.io) library was used to develop the web interface that provides dynamic updates based on user inputs in the graphical user interface (GUI) without writing or editing code directly. Streamlit was chosen due to its simplicity in development and compatibility with other Python libraries across operating systems. Streamlit v1.39.0 is compatible with any modern HTML5 web browser. For data loading and processing, we relied on Pandas v2.2.3 with PyArrow v18.0.0 which achieves faster data loading and processing compared to Pandas alone. We used NumPy v1.26.4 for data type and range selection, RGB array creation for color handling, and grid setup for subplots. Seaborn v0.13.2, Matplotlib v3.9.2, and Plotly v4.24.1 libraries were used for data visualization and users are provided with options to adjust parameters: plot size, color profile, and output formats in PNG, JPG, SVG, or PDF. CAFE integrates AnnData^12^, a widely used framework in single-cell RNA sequencing analysis that allows for efficient storage and manipulation of both sparse and dense matrices along with metadata. CAFE outputs an AnnData object which can be used outside of CAFE if users wish to deposit their data with analysis or perform custom analyses using other tools.

**Figure 1:**
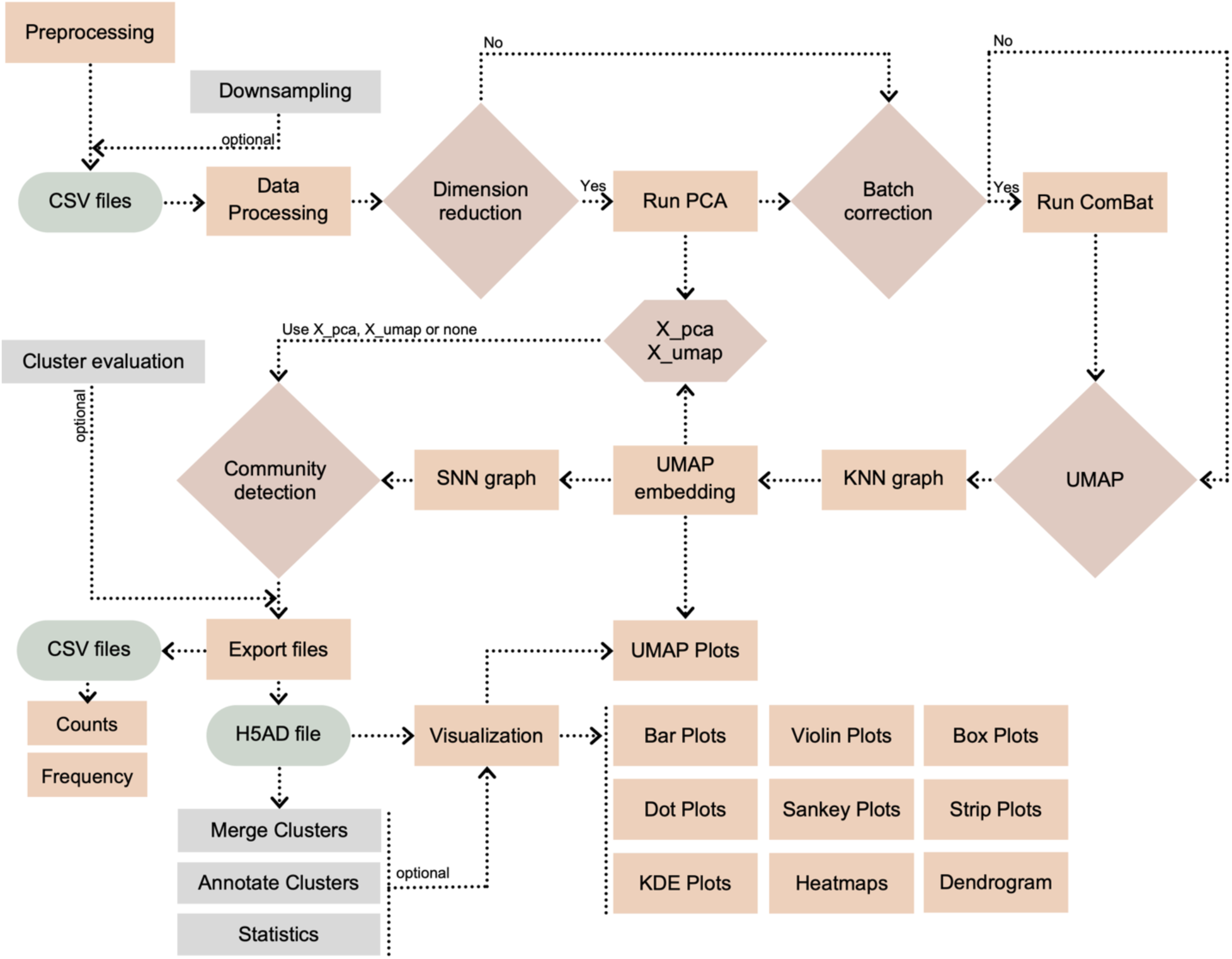
The flowchart outlines steps and components of CAFE’s workflow. Preprocessing includes compensation, data scaling/transformation using a standard FCM software and scaled CSV files are then exported and renamed as Sample_Group.csv. Data processing performs error checks and concatenation of CSV files into an AnnData object/H5AD file. Major steps requiring user input include dimension reduction, batch correction, UMAP (UMAP Uniform Manifold Approximation and Projection) and community detection. Outputs are downloadable as CSV, H5AD, PNG, JPG, SVG and PDF files.

The Scanpy library was used to perform key analyses including dimension reduction, batch correction, and Leiden clustering. The user has the option to reduce dimension (sc.tl.pca) of the data through Principal Component Analysis (PCA) or skip it. PCA is a linear dimensionality reduction technique that retains the global structure of the data by capturing the variances across all dimensions. Because the app performs PCA through Scanpy library, by default, the number of components retained is limited to the lesser of the two values: the number of cells or the number of markers. Also, the Singular Value Decomposition (SVD) solver was set to “auto” that chooses the most appropriate solver based on the size of the dataset; however, users have options to set a percentage of variances they want to retain, and the type of solver used. The reduced dataset is stored and can be further processed for batch correction (sc.pp.combat) using ComBat (Combined Batch)^17^. This is particularly useful if a user has collected samples in different batches as the algorithm standardizes the data by making it comparable and removing unwanted variability.

To group cells into distinct clusters based on marker expression profiles, Leiden clustering is run (sc.tl.leiden) and users can select either ‘iGraph’ or ‘leidenalg’ algorithm flavor^10,18^. To define clustering resolution, a user can choose from 0.01 to 2.0 where the lowest value provides the lowest number of clusters. The user can fine tune Leiden calculation by altering the number of neighbors and minimum distance values in Uniform Manifold Approximation and Projection (UMAP) calculation within the app. CAFE generates AnnData object (H5AD) file, CSV outputs, and visual outputs including UMAP plots, dot-heatmaps, expression pots, and barplots as high-resolution images and provides download buttons to save them to a desired folder. The app allows various visualization settings, with changes made and displayed immediately within the app. The generated AnnData object can be further used to perform a range of statistical analyses.

The app includes advanced functionalities for clustering and cluster evaluation. Because setting up appropriate values for Leiden resolution and UMAP parameters is central to obtaining quality clustering results, a user can leverage CAFE’s Cluster Evaluation tab to generate multiple AnnData files with various combinations of these parameters and compare UMAP plots as well as Silhouette score, Calinski-Harabasz score, Davies-Bouldin score, and Elbow method to assess clustering results. Besides, CAFE provides clustering with pre-selected markers, merging subclusters into metaclusters, and annotation of clusters directly within the app. The advanced Downsampling tab offers to downsample (e.g. 20,000 events per sample) data using a PCA-KDE based method. This approach combines PCA and KDE (Kernel Density Estimation) based algorithm from scipy.stats using Gaussian kernel function and silverman bandwidth^19^. KDE is applied to the PCA-transformed data to estimate the density of data points. Based on these density estimates, the code probabilistically downsamples data, thus reducing sampling bias and preserving original data distribution. This method offers an informed approach compared to simple random downsampling, and it can be used to filter out noise while retaining meaningful biological information in a smaller dataset.

### Statistical analysis

In the visualization tab under statistical analyses section, users can perform different statistical tests and generate plots. The Shapiro-Wilk test from “scipy.stats” is used to determine if marker expression within each group or cluster follows a normal distribution. Based on these results, the app recommends either parametric (T-test) or non-parametric (Mann-Whitney U test) tests using “scipy.stats”. For comparing multiple clusters, the app allows users to perform ANOVA (scipy.stats.f_oneway) or Kruskal-Wallis (scipy.stats.kruskal) tests. To reduce statistical artifacts, multiple testing correction is applied using the Benjamini-Hochberg False Discovery Rate (FDR) through “statsmodels.stats.multitest”. Additionally, effect size measures are computed to complement statistical p-values, with users able to choose between parametric tests (Cohen’s d) or non-parametric tests (Cliff’s Delta). Cohen’s d is calculated using basic functions from Numpy, while Cliff’s Delta is computed with the “cliffs_delta” package. To assess associations between clusters and groups, we used Chi-square testing from “scipy.stats.chi2_contingency” and contingency tables with “pandas.crosstab”. Residual calculations were displayed in Streamlit as tables to help users understand which clusters are more prevalent within certain groups.

### Performance and reproducibility

We have set a global setting for Scanpy (sc.settings.n_jobs = −1) to use all available CPU cores. For advanced clustering, multi-threading was achieved using Python’s joblib library. Two other libraries were used, watchdog v5.0.2 and iGraph v0.10.8^18^. Watchdog helps in monitoring file change events and improves performance of Streamlit by providing real-time feedback. iGraph is designed to handle complex networks and graph operations and is used by Scanpy as part of the Leiden clustering to perform graph operations. We recommend ‘iGraph’ over ‘leidenalg’ as iGraph is implemented in C and achieves advantages in performance compared to high-level interpreted languages such as Python. To export Plotly figures, we have used Kaleido engine v0.2.1. We have tested the app with various datasets using an Apple M3 Pro System with 18GB of random-access memory (RAM). CAFE is primarily intended to be used using local computer; however, it can be scaled up using any High-Performance Computing (HPC) system that supports an HTML5 web browser. We also provided scripts in our GitHub page to generate AnnData with dimension reduction and Leiden clustering through command-line interface (CLI) based HPC systems.

## Results

To demonstrate the functionality of the app, we have analyzed 35-color spectral flow cytometry data (Publicly available at FlowRepository: FR-FCM-Z3WR) of human peripheral blood mononuclear cells (PBMC) obtained from COVID-19 hospitalized patients and healthy controls^20^. A total of 10 samples were analyzed with 5 from each group. For best practices, we installed and ran CAFE through Pixi package manager. Users can also install and run the app using Anaconda package manager as described in our Github documentation. Once initiated through a terminal (*pixi run cafe, or python cafe.py*), a web browser opens with the CAFE app at localhost on port 8501. The default data loading limit is set to 3GB, but a user can change the value from the cafe.py script if necessary.

### Data processing

The uploaded public data were available as doublets-debris removed and CD45+ gated; so we obtained the CSV files just by exporting scaled values from FlowJo v10.10.0. Data scaling is generally recommended for high resolution clustering but there may be instances where users may use raw values. Data can be similarly exported from other flow cytometry software such as FCS Express. It is required that flow cytometry data have proper compensation. We recommend manual inspection of flow cytometry data and removal of debris, dead cells, and doublets prior to exporting the scaled files. A user can also gate on appropriate cell type and export the data to obtain more focused clustering results. To streamline downstream analysis, we have implemented a naming convention for the CSV files. Each CSV file name must begin with a unique “SampleName” followed by “GroupName”, separated by an underscore; for instance, “Sample01_Control.csv” and “Sample02_Treatment.csv”. After loading the data, the app will import the required libraries and perform initial checks for data structure and incorporate SampleID and GroupName into the dataframe based on the CSV file names. Within the dataframe, rows containing any missing values are skipped and anomalies in data structure are reported. In this study, we used the advanced KDE-based downsampling option in CAFE to downsample data to 35,000 cells per sample for a total of 350,000 cells and 12.25M data points (number of cells multiplied by number of markers). This is an optional step prior to data processing. After loading the files, the app processed (7.8 sec) and combined the expression data and metadata without errors to create an AnnData object.

### Dimension reduction and batch effects

After generating the AnnData object (H5AD file), we selected dimension reduction using PCA with SVD solver set to auto and retained components with 95% variance. The app ran PCA (2.16 sec), kept 12 components and generated (PC1 x PC2) graphs by Groups. Depending on the data size, a user can choose from auto, full, arpack, and randomized SVD solver. Randomized, for example, is better suited for larger datasets as it provides a balance between speed and accuracy. For batch correction, we applied ComBat (1.06 sec) and proceeded to Leiden clustering.

### Leiden clustering and metaclustering

For the dataset, we applied Leiden resolution of 1.0 with flavor set to iGraph, UMAP n_neighbors to 15, min_dist to 0.1 and distance calculation method as Euclidean. A user has the option to use a slider control to choose from resolution values 0.01 to 2.0. To find the optimal resolution, we initially made use of Advanced Cluster Evaluation option in CAFE to generate a series of AnnData files with varied Leiden resolution and n_neighbor values and observed the UMAPs to find distinct clusters that are biologically meaningful for the dataset. With Leiden resolution of 1.0, we initially obtained a total of 30 clusters for the PBMC dataset which took 11.5 minutes for calculation. Once clustering was completed, CAFE generated a frequency table of each sample by Leiden cluster for the number of cells, frequency of cells, and median fluorescence intensity of each marker for each cluster. Using these 3 tables, users can perform statistical analyses to compare cluster count and frequency by groups and expression of marker proteins within clusters by group. Using the Advanced Cluster Merging option, we merged the subclusters with similar profile into corresponding metaclusters resulting in a new total of 16 clusters.

### Characterization of PBMC Subpopulations

To characterize the phenotypic properties, we examined the expression of surface markers for each identified cluster by protein expression UMAP plots (Figure 2a). We found high CD3 expression in T cells clusters with CD4 and CD8 expression showing corresponding T cell subtypes. CD8+ effector memory (Tem) and central memory (Tcm) subsets were differentiated by high CCR7 and CD27 expression in Tcm. We used CD45RA expression to identify terminally differentiated Tem cells (Temra). Monocyte clusters were identified by CD14 and CD16 expression, distinguishing classical monocytes (cMO) from other monocyte subsets, while natural killer (NK) cells showed high levels of CD56, corresponding with CD56^Bright^ NK cells. Based on shared marker profiles and hierarchical ranking, T cell subsets (Tem, Tcm, and Temra) formed a distinct grouping separate from B cells and myeloid-derived cells, reflecting the differential expression of lineage-specific markers.

**Figure 2:**
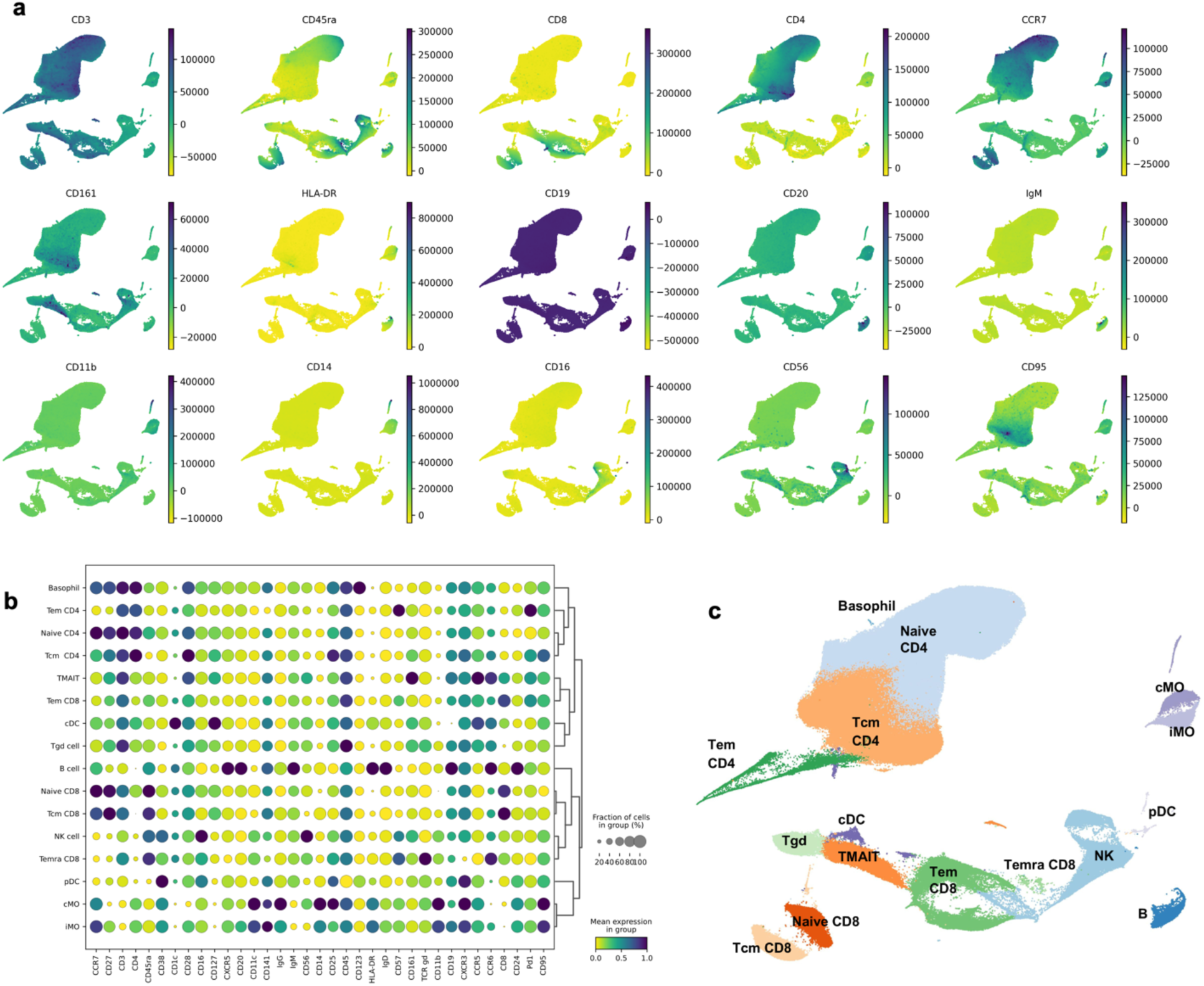
Profiling of Human PBMCs Reveals Distinct Immune Subpopulations and Marker Expression Patterns. (**a**) UMAP plots showing selected marker expression intensities across all cells in the UMAP space to highlight lineage-specific marker distribution. (**b**) Dot plot of all marker expression across all identified PBMC cell types. Dendrogram highlighted distinct marker-based groupings. (**c**) UMAP visualization showing 16 distinct clusters with annotated cell types including Naive CD4 and CD8 T cells, central memory CD4 and CD8 T cells (Tcm), effector memory CD8 T cells (Tem), terminally differentiated effector memory CD8 T cells (Temra), mucosal-associated invariant (MAIT) T cells, classical monocytes (cMO), intermediate monocytes (iMO), B cells, NK cells, gamma delta (γδ) T cells (Tgd), conventional dendritic cell (cDC) and plasmacytoid dendritic cell (pDC).

Based on the expression profiles of marker proteins, we annotated the clusters using CAFE’s Advanced Annotation tab and classified them into 16 distinct cell types. We also used the dotplot to confirm annotations of the cell types (Figure 2b). For instance, the B cell-specific marker CD19 and CD20 were used to identify the B cell cluster, the CD14 marker to identify monocytes, and the CD16 marker to identify NK cells. High CD20 expression in B cell cluster indicated their mature stage in immune response. Our annotated UMAP (Figure 2c) shows well-defined clusters that correspond to PBMC lineages, including Naive CD4+ and CD8+ T cell and Tcm for both CD4+ and CD8+ subsets. We also identified Tem and Temra cells, as well as mucosal associated invariant CD8+ T cell (MAIT). We also identified cMO, intermediate monocytes (iMO), B cells, NK cells, Gamma delta (γδ) T cells (Tgd) and dendritic cell (DC) types (cDC and pDC).

### Distinct Cellular and Molecular Signatures Observed in COVID-19 Compared to Healthy Controls

UMAP analysis of COVID-19 hospitalized patients compared to healthy controls revealed distinct clustering patterns between groups, particularly among monocytes, NK cells, and CD8 T cells (Figure 3a). To understand changes between the two groups, CAFE offers varied visualization options, for instance, we used a Sankey diagram to demonstrate that MAIT cells and Tgd are much less abundant in COVID-19 patients compared to healthy controls (Figure 3b). We also found that CD8 Tcm and B cells were significantly expanded in COVID-19 patients. A composite bar-strip plot also demonstrates the distribution of cells in frequency where each dot represented each sample colored by specific group (Figure 3c). The total number of cells in iMO were largely reduced in COVID-19 patients compared to healthy controls (Figure 3d). These data may indicate a possible shift from an innate response towards an adaptive response. To quantify the effect size of changes observed, we compared cell types within the COVID-19 group to healthy controls as a reference and found changes in naive CD8, TMAIT, and iMO cells have a larger effect size, demonstrating a bigger difference between the two groups (Figure 3e). We further compared these cell types by plotting box plots for individual cell types (Figure 3f) which demonstrated a non-statistically significant increase in Tcm CD8 (p=0.0777) and statistically significant decrease in naive CD8 cells (p=0.0372) in COVID-19 patients compared to healthy controls.

**Figure 3:**
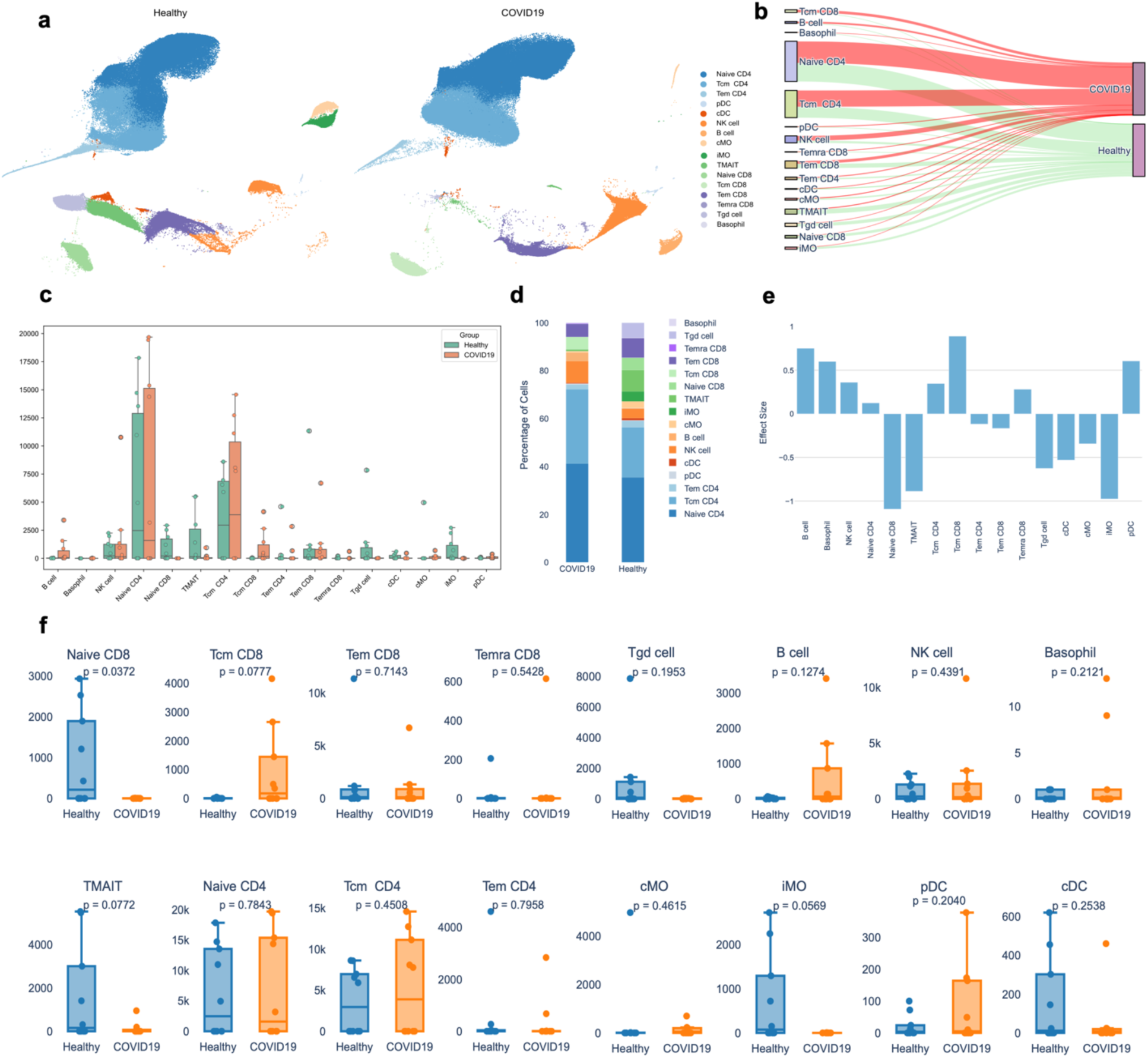
Comparative Analysis of Immune Cell Subpopulations in Healthy and COVID-19 Individuals. (**a**) UMAP plots displaying distinct clustering patterns and differential distribution of cell types in PBMC across healthy and COVID-19 group. (**b**) Sankey diagram illustrates the distribution of cells across groups, with thicker flow indicating more cells. (**c**) Composite bar-strip plot summarizing cell count distribution across cell subpopulations. Dots represent each individual samples colored by group. (**d**) Stacked bar chat showing distribution of cells in percentage across two groups. (**e**) Effect size calculated using Cohen’s d indicating changes in the number of cells in COVID-19 compared to reference healthy control. (**f**) Comparison of individual cell type frequencies between healthy and COVID-19 groups with p-values for statistical significance. N=9/group.

### Altered Marker Expression Profiles in COVID-19 Patients

CAFE outputs a file with expression data for all markers on each cluster by sample for use in external plotting and statistical software. In addition, further exploration of specific markers within clusters can be performed within CAFE. We used this approach to identify that MAIT cells have the greatest expression of CD161, as shown by violin plot (Figure 4a). We also examined marker distribution in CD8 T cell populations and found that more cells within the Tem CD8 population appeared activated based on greater HLA-DR, CXCR3, and CCR5 expression compared to other CD8 T cell subsets (Figure 4b). We found that median expression of CD8, CD14, CD11b, IgG were increased in COVID-19 patients compared to healthy controls across all clusters (Figure 4c). These reflect the overall differences in some of the cell populations we observed between groups. We examined marker expression within the B cell cluster and found that more B cells expressed IgG in COVID-19 samples compared to healthy controls although the difference was not statistically significant (p=0.1631), while IgD expression was statistically significantly reduced (p=0.0162) in COVID-19 samples compared to healthy controls (Figure 4d). We also examined activation markers in CD8+ T cells and found that COVID-19 patients had more CD8+ T cells that expressed CD25 and HLA-DR than healthy controls (Figure 4e).

**Figure 4:**
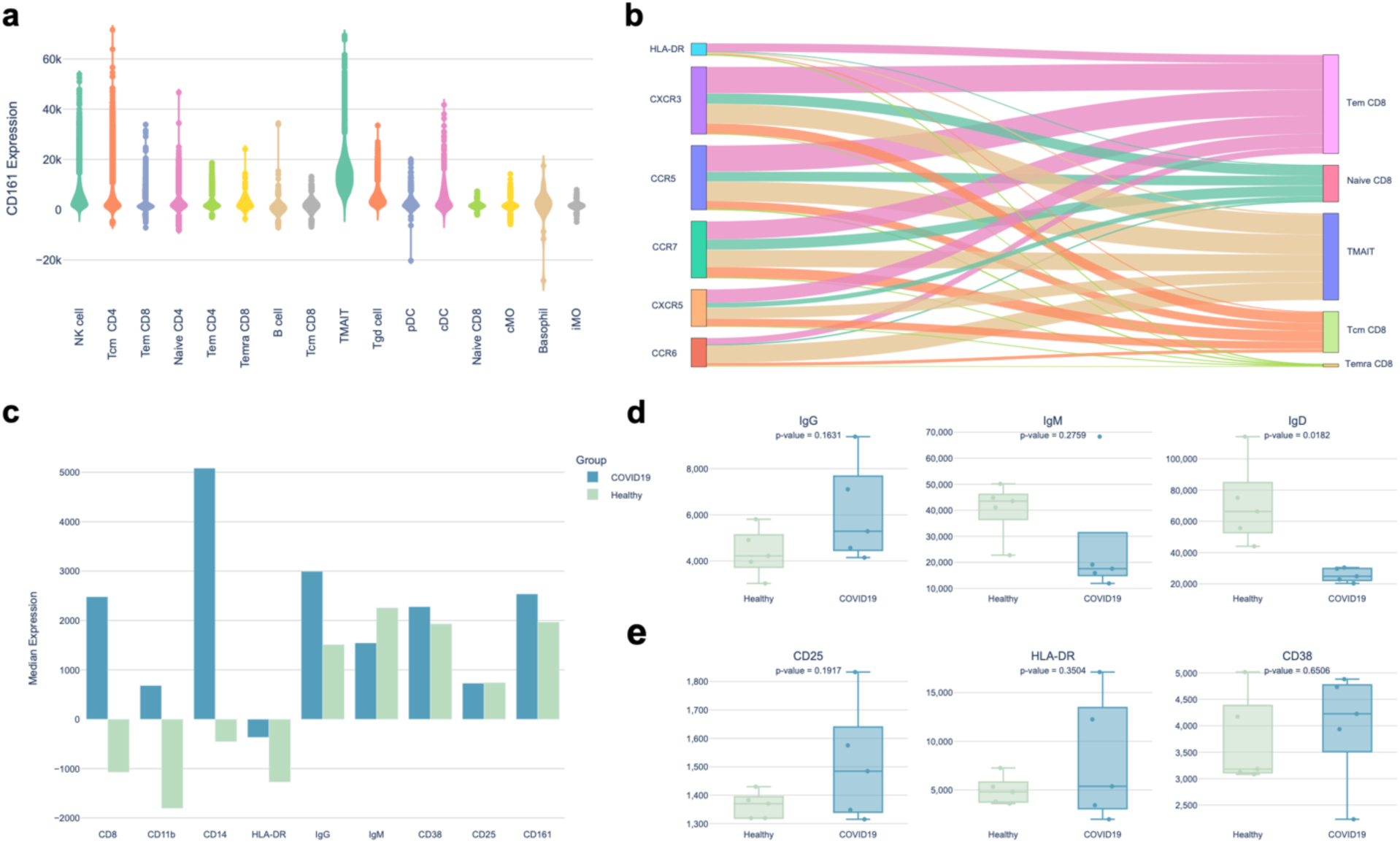
Marker expression and distribution differences between COVID-19 and healthy individuals. (**a**) Violin plot showed the median expression levels of CD161 across immune cell subtypes. (**b**) Sankey diagram illustrated marker expression in CD8+ T cells, with thicker flows indicating more cells expressing that marker. (**c**) Bar chart showed median expression of markers across all cell types between COVID-19 and healthy individuals where positive values indicated upregulation. Box plots displayed (**d**) the number of cells expressing IgG, IgM and IgD in B cells, and (**e**) the number of CD8+ T cells expressing CD25, HLA-DR, and CD38.

## Discussion

As research in immunology increasingly relies on high-dimensional cytometric data, there is a growing need for a user-friendly analysis tools for everyday use. Here, we present CAFE as a free and open-source tool designed to address the analytical and accessibility issues posed by SFCM data. CAFE uses a GUI and interactive controls to enable immunologists to analyze complex data without needing specialized coding knowledge.

Using a jupyter notebook, we have previously shown the ability of Scanpy’s Leiden function (*scanpy.tl.leiden*) to analyze a 50-color human PBMC dataset^15^. CAFE acts as a wrapper combining packages within Streamlit to provide a web interface, offering more accessible and extensive functionality than Jupyter notebooks’ CLI. We demonstrated analysis of a 35-color human PBMC dataset using CAFE in this study with 350,000 cells and 12.25M data points. Major steps including data processing, dimension reduction, batch correction and Leiden clustering were completed in under 12 minutes using an Apple M3 Pro laptop. In our analysis, we observed COVID-19 patients with altered immune cell distributions and marker expression profiles, consistent with prior findings, as well as MAIT cells expressing high CD161 and B cells expressing high CD20^20^.

While developing CAFE, we have balanced compatibility and performance and included many options for customization of how the code processes and analyzes data while integrating default options and tooltips to help guide users. Our implementation of Pandas with PyArrow significantly improves processing speed over Pandas alone. However, transitioning to Polars’ lazy evaluation framework could further speed up processing once compatibility issues between ARM and x86 machines are resolved. CAFE’s web app design and functionality also revolved around simplicity as we de-emphasized features that are not commonly used in order to streamline user-experience. For data dimension reduction, we adhered to a convention of using PCA as the primary method and using PCA-reduced representations for constructing UMAP neighbor graphs, as opposed to utilizing UMAP directly for primary dimensionality reduction. Although PCA is designed for linear data, it effectively reduces noise and enhances clustering performance. Users have the choice to skip PCA to perform Leiden clustering on the raw data or use UMAP embeddings (i.e., X_umap) to use UMAP reduced data for clustering. One viable alternative to PCA for non-linear data structure is Kernel PCA, but we have skipped adding the kernel PCA option in the CAFE workflow because it may not be practical since it is computationally taxing.

UMAP parameters and Leiden resolution largely influence the number of clusters for community detection. Leiden clustering is performed on the graph structure, so evaluation of clustering quality solely based on methods such as elbow or silhouette score is not ideal. Rather a combined approach with prior biological knowledge can inform the most correct clustering resolution. We recommend using CAFE’s advanced clustering evaluation tab to generate plots with a range of varied UMAP parameters and Leiden resolutions for visual inspection. Using this approach, users can select the most appropriate clustering resolution for each dataset. Since this is an unsupervised algorithm, setting up an incorrect resolution can heavily skew the interpretation of data.

Manual gating continues to be the gold standard in flow cytometry analysis, but it is limited by sequentially drilling down into subsets of cells with 2-dimensional bi-axial gating. Thus, our goal was to complement this hypothesis-driven approach with the unsupervised computational algorithms. In this way, users can perform hypothesis-driven analysis with manual gating and hypothesis-generating analysis with unsupervised clustering. Compared to other open-source tools, CAFE provides a wide range of publication-ready visualization options. Due to its underlying code in python, CAFE is highly scalable to datasets of millions of cells and takes advantage of multi-threading to obtain higher performance. Previously, python-based Pytometry and CRUSTY integrated unbiased clustering algorithms within their tools^21,22^. Pytometry incorporates the Leiden algorithm^10^, which has been shown to be an improvement over the predecessor Louvain algorithm^23^; however, Pytometry requires coding using Python. CRUSTY incorporates an easy-to-use GUI but it does not offer the Leiden algorithm and relies on FlowSOM and Phenograph. Another limitation of CRUSTY is that the cloud-based service limits users to analyzing 100,000 total events. There are also limited visualization and analysis options in CRUSTY and they rely on most of the Phenograph and FlowSOM default settings which cannot be customized. Cloud-based solutions may face limitations in availability, scalability, and data security. Users may be prohibited from uploading data to cloud-based systems that have protected health information due to HIPAA. Among a few other GUI based tools, Cytoflow^24^, Floreada (floreada.io), EasyFlow^25^ allow for flow cytometry data analysis but do not offer clustering. FlowPy (flowpy.wikidot.com) allows for clustering but uses k-mean clustering rather than the most advanced algorithms currently in use (i.e. Leiden). Additional tools like terraFlow^26^ and CellEngine (CellCarta, Montreal, Canada) are for-profit spectral flow cytometry analysis softwares and the price of these may be restrictive for many users. Finally, FlowJo is a staple for many immunologists and has some native clustering capabilities. It also supports plugins for additional clustering algorithms, but these add-ons do not offer much customization in the clustering parameters.

CAFE, while addressing many of these limitations as an open-source alternative, has its own practical considerations. CAFE is intended to be run locally which requires installing a package manager such as Pixi or Anaconda/Miniconda3 through terminal. Performance is also dependent on the user’s machine. For larger datasets, we recommend utilizing our provided scripts in Github to run data processing step through an HPC cluster by allocating more RAM. Once the user has Anndata file generated with cluster information, all the downstream analysis and figure generation steps become significantly less computationally demanding. Ultimately, CAFE’s aim is to become a secure, scalable, and open-source platform accessible to a broad range of researchers to run complex analyses through a simple intuitive graphical user interface.

## Acknowledgements

This work was supported by the National Institutes of Health/National Institute on Aging Grant R00 AG068309 (to D.J.T.); Nathan Shock Center, which is supported by the National Institutes of Health/National Institute on Aging Grant P30 AG050886; and this research was conducted while (Daniel Tyrrell) was an AFAR Grant for Junior Faculty awardee A24063 (to D.J.T.).

## Data availability

The raw data are publicly available at FlowRepository: FR-FCM-Z3WR. The downsampled .csv files are available from Figshare: 27940719. The processed Anndata object after dimensionality reduction and clustering is also available from Figshare: 27940752.

## Code availability

CAFE is freely available at https://github.com/mhbsiam/cafe.

## Contributions

M.H.B.S. and D.J.T. conceived the software development. M.H.B.S. created the software and web application; M.H.B.S. analyzed the data. M.H.B.S. and D.J.T. wrote the paper. All authors reviewed and edited the paper.

## Ethics declarations

The authors declare they have no competing interests.

## Notes

http://flowrepository.org/id/FR-FCM-Z3WR

https://figshare.com/articles/dataset/Downsampled_data_from_FlowRepository_FR-FCM-Z3WR/27940719

https://figshare.com/articles/dataset/Processed_h5ad_Anndata_file_from_Downsampled_data_to_use_for_Visualization_/27940752

https://github.com/mhbsiam/cafe

### Competing Interest Statement

The authors have declared no competing interest.

### Summary of Updates

Corrected some typos in the manuscript throughout all sections.

## References

1. McKinnon, K. M. Flow Cytometry: An Overview. Current Protocols in Immunology 120, 5.1.1–5.1.11 (2018).

2. Nolan, J. P. The evolution of spectral flow cytometry. Cytometry Part A 101, 812–817 (2022).

3. Konecny, A. J., Mage, P. L., Tyznik, A. J., Prlic, M. & Mair, F. OMIP-102: 50-color phenotyping of the human immune system with in-depth assessment of T cells and dendritic cells. Cytometry Part A 105, 430–436 (2024).

4. Tyrrell, D. J. et al. Clonally expanded memory CD8+ T cells accumulate in atherosclerotic plaques and are pro-atherogenic in aged mice. Nat Aging 3, 1576–1590 (2023).

5. Na, S., Choo, Y., Yoon, T. H. & Paek, E. CyGate Provides a Robust Solution for Automatic Gating of Single Cell Cytometry Data. Anal Chem 95, 16918–16926 (2023).

6. Van Gassen, S. et al. FlowSOM: Using self-organizing maps for visualization and interpretation of cytometry data. Cytometry A 87, 636–645 (2015).

7. Samusik, N., Good, Z., Spitzer, M. H., Davis, K. L. & Nolan, G. P. Automated mapping of phenotype space with single-cell data. Nat Methods 13, 493–496 (2016).

8. Qiu, P. et al. Extracting a cellular hierarchy from high-dimensional cytometry data with SPADE. Nat Biotechnol 29, 886–891 (2011).

9. Levine, J. H. et al. Data-Driven Phenotypic Dissection of AML Reveals Progenitor-like Cells that Correlate with Prognosis. Cell 162, 184–197 (2015).

10. Traag, V. A., Waltman, L. & van Eck, N. J. From Louvain to Leiden: guaranteeing well-connected communities. Sci Rep 9, 5233 (2019).

11. Hao, Y. et al. Integrated analysis of multimodal single-cell data. Cell 184, 3573–3587.e29 (2021).

12. Wolf, F. A., Angerer, P. & Theis, F. J. SCANPY: large-scale single-cell gene expression data analysis. Genome Biology 19, 15 (2018).

13. Rich, J. M. et al. The impact of package selection and versioning on single-cell RNA-seq analysis. bioRxiv 2024.04.04.588111 (2024) doi:10.1101/2024.04.04.588111.

14. McInnes, L., Healy, J., Saul, N. & Großberger, L. UMAP: Uniform Manifold Approximation and Projection. Journal of Open Source Software 3, 861 (2018).

15. Vardaman, D. et al. Development of a Spectral Flow Cytometry Analysis Pipeline for High-dimensional Immune Cell Characterization. J Immunol 213, 1713–1724 (2024).

16. Burton, R. J. et al. CytoPy: An autonomous cytometry analysis framework. PLoS Comput Biol 17, e1009071 (2021).

17. Johnson, W. E., Li, C. & Rabinovic, A. Adjusting batch effects in microarray expression data using empirical Bayes methods. Biostatistics 8, 118–127 (2007).

18. Csardi, G. & Nepusz, T. The Igraph Software Package for Complex Network Research. InterJournal Complex Systems, 1695 (2005).

19. Silverman, B. W. Density Estimation for Statistics and Data Analysis. (Routledge, New York, 2017). doi:10.1201/9781315140919.

20. Yu, C. et al. Mucosal-associated invariant T cell responses differ by sex in COVID-19. Med 2, 755–772.e5 (2021).

21. Büttner, M., Hempel, F., Ryborz, T., Theis, F. J. & Schultze, J. L. Pytometry: Flow and mass cytometry analytics in Python. 2022.10.10.511546 Preprint at 10.1101/2022.10.10.511546 (2022).

22. Puccio, S. et al. CRUSTY: a versatile web platform for the rapid analysis and visualization of high-dimensional flow cytometry data. Nat Commun 14, 5102 (2023).

23. Blondel, V. D., Guillaume, J.-L., Lambiotte, R. & Lefebvre, E. Fast unfolding of communities in large networks. J. Stat. Mech. 2008, P10008 (2008).

24. Teague, B. Cytoflow: A Python Toolbox for Flow Cytometry. 2022.07.22.501078 Preprint at 10.1101/2022.07.22.501078 (2022).

25. Ma, Y., Eizenberg-Magar, I. & Antebi, Y. EasyFlow: An open-source, user-friendly cytometry analyzer with graphic user interface (GUI). PLOS ONE 19, e0308873 (2024).

26. Freeman, D., et al. terraFlow, a high-parameter analysis tool, reveals T cell exhaustion and dysfunctional cytokine production in classical Hodgkin’s lymphoma. Cell Reports 43, (2024).

